# Using Cryogenic Electron Tomography (cryoET) to Determine Rubisco Polymerization Constants in α-Carboxysomes

**DOI:** 10.64898/2026.03.20.713215

**Authors:** Wenxiang Cao, Kristy Rochon, Ryan H. Gray, Luke Oltrogge, David F. Savage, Enrique M. De La Cruz, Lauren Ann Metskas

## Abstract

Bacteria microcompartments (BMCs) are pseudo-organelles comprised of a self-assembling, semi-permeable protein shell, most commonly enclosing components of enzymatic pathways. α-Carboxysomes (α-CBs) are anabolic BMCs known for their role in sequestering Rubisco, the enzyme responsible for carbon fixation in plants, algae and bacteria, along with an upstream enzyme and an assembly protein. Rubisco has low selectivity for its substrate, CO_2_, and has a slow enzymatic turnover rate, resulting in an inefficient metabolic pathway. Within the α-CB, Rubisco has been observed at a range of concentrations and with either a liquid-like assembly or a pseudo-lattice of polymerized fibrils. The biophysical origins of the fibril ultrastructure organization are unclear; however, it is only observed inside α-CBs. Quantitative knowledge of the binding constants and energies for assembly and maintenance of these fibrils is critical for understanding this organization and Rubisco regulation, but quantitative methods for *in situ* analysis of Rubisco polymerization have been lacking. Here, we present an approach to convert tomography-derived α-CB volumes and Rubisco particle positions into polymerization binding curves. We used this procedure to determine the Rubisco polymerization constants, including the nucleus size (*n*) and equilibrium polymerization constant (*K*_*pol*_). The adopted modeling approach is consistent with *in situ* constraints, such as concentration-dependent binding interactions and confinement. This approach offers a powerful tool to evaluate both *in vitro* and potentially *in vivo* biomolecular interactions, both of Rubisco and of other proteins and polymers suitable for analysis by cryo-electron tomography.

**Significance Statement:** Cryogenic electron tomography (cryoET) is a powerful method to resolve structures of proteins in their native environment at subnanometer-level resolution. Because tomography data retains spatial relationships of all particles, it intrinsically contains information about component (e.g., protein) binding interactions. Here, we use Rubisco polymerization in α-carboxysomes as a model system to demonstrate that quantitative, biochemical binding analysis is possible with cryoET.

## Introduction

Quantitative knowledge of the equilibrium binding affinities of (macro) molecular interactions is crucial for understanding the regulation of key biological processes including enzyme catalysis, signal transduction, gene expression, and the assembly of macromolecular complexes (1–4). Established methods for determining molecular binding constants commonly rely on experimental data acquired *in vitro*, under controlled experimental conditions (5, 6). Such *in vitro* studies are indispensable for developing quantitative, mechanistic models, but they often have inherent limitations, such as restricted concentration ranges, non-native chemical modifications (i.e., tags), and/or minimal buffer systems substituting for uncharacterized physiological chemical environments. Studies rarely evaluate the effects of physiological crowding (7, 8), and thus fail to replicate native physiological conditions.

Cryogenic electron tomography (cryoET) is a powerful imaging technique with resolutions ranging from near-atomic (protein structures) to micron (cellular features) (9–11). While the method typically provides slightly lower resolution than cryogenic electron microscopy (cryoEM) in structural biology applications, it has the unique ability to preserve the spatial relationships between imaged particles. In subtomogram averaging (STA), proteins are identified, extracted from the larger image, aligned and averaged, and then replaced into the tomogram, enabling extrapolation of these spatial relationships (12, 13). Despite these capabilities, the most detailed analyses of proteins with cryoET typically focus on structure, with more qualitative or morphological descriptions of protein environment, proximity to other features, and ultrastructure arrangement.

Here, we present a method using cryoET to quantify binding interactions and polymerization constants of a self-associating biopolymer *in situ*. We posit that if all particles in a defined and precisely measured volume are accurately identified and their oligomeric states distinguished, biophysical binding parameters such as polymerization constants can be calculated from STA data. In addition to providing simultaneous insight into protein conformation and binding, this approach eliminates the need for complex *in vitro* assay design while maximizing the data available in tomograms and extending polymerization assay capabilities to systems not amenable to *in vitro* measurements.

We use Rubisco in α-carboxysomes (α-CBs), a bacterial microcompartment (BMC) responsible for carbon fixation in *Halothiobacillus neapolitanus* (14–16), as a model system to validate this approach. α-CBs are ∼1 aL pseudo-organelles with a selectively permeable protein shell (17–20). The protein shell assembles around an enzyme condensate, facilitated by a disordered scaffolding protein (21). α-CBs encapsulate Rubisco with its upstream enzyme carbonic anhydrase, maximizing Rubisco turnover rates presumably through substrate saturation (14, 19, 22–24). α-CB assembly and cargo sequestration rely largely on Rubisco condensation through liquid-liquid phase separation (25). This process requires a disordered accessory protein (CsoS2) interfacing with the shell to assemble the facets while also forming interfaces with Rubisco to facilitate inclusion within the microcompartment (21). The shell is assembled around the condensate.

Rubisco condensates with CsoS2 inside α-CBs assemble pseudo-lattices of polymer fibrils, a behavior not replicated with purified components *in vitro* (Fig. 1A) (21, 26, 27). Recent studies have indicated that BMC shells can separate redox states and that the interior of β-CBs becomes oxidized over time, with unknown implications on the BMC’s interior organization or function (28, 29).

**Figure 1.**
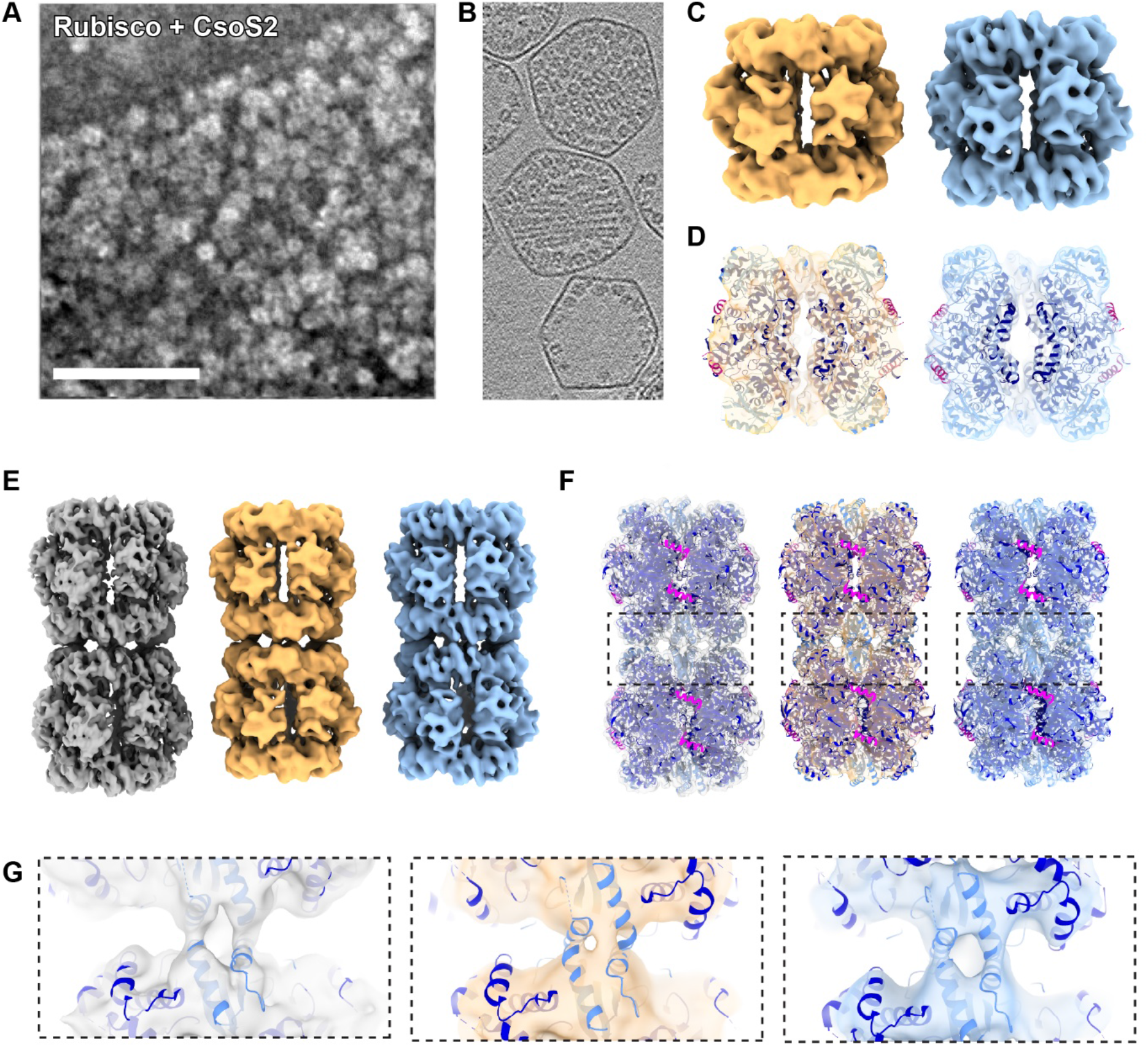
Aligned Rubisco Particles from Tomograms. **A**. Negative stain TEM micrograph of a Rubisco-CsoS2 condensate. **B**. Orthoslice of a tomogram containing α-CBs with examples of Rubisco packing: dense (top), ordered (middle), and sparse (bottom). **C**. Rubisco densities resulting from STA of DTT treated (left, orange) and diamide treated (right, blue) samples. **D**. Docked crystal structure (PBD ID: 6UEW) in DTT-treated density (left, yellow) and diamide-treated density (right, blue). **E-G**. Composite chains of STA results with docked crystal structures of (PDB ID: 6UEW) for untreated Rubisco (left, gray), DTT-treated (middle, orange) and diamide-treated (right, blue). **G**. Isolated Rubisco-Rubisco interfaces resolved using STA. Untreated (left, gray), DTT-treated (middle, orange) and diamide-treated (right, blue).

In this study, we developed and validated a framework and procedure to quantitatively extract polymerization binding constants directly from cryoET data, using Rubisco in α-CBs as a model system. By converting STA volumes and particle positions into polymerization “binding” curves, we determined key binding equilibrium constants governing Rubisco fibril assembly *in situ*. This approach not only addresses a critical gap in our ability to quantify biomolecular assembly under native physiological conditions, offers a complementary technique to traditional *in vitro* methods and extending the analytical applications of cryoET beyond structural studies, but also provides a general method in analysis of biomolecule polymerization with the theories already developed.

## Results

### Rubisco oligomeric states within α-CBs

Rubisco within α-CBs can adopt two oligomeric/polymer states (Fig. 1B): dense (disorganized or liquid-like) and ordered (lattice-like packing). Purified α-CBs can also have low Rubisco concentration (termed “sparse”) as an experimental consequence of centrifugation (26). It is not known what determines whether Rubisco within α-CBs adopts the dense or ordered morphology at equivalent concentrations. We hypothesize that Rubisco polymerization could be affected by redox state. To test this, α-CBs were isolated from *H. neapolitanus* and treated with either DTT (reducing agent) or diamide (oxidizing agent) to alter their internal redox state before vitrification and cryoET analysis of polymerization.

Rubisco monomers, oligomers, and linear polymers were identified within intact α-CBs and STA performed out to sub-7Å resolution using published methods (Fig. 1C) (26, 30). Helices are well defined at this resolution, allowing precise estimation of position and orientation for each Rubisco complex (Fig. 1D). We used a numerical classification approach to calculate α-CB volume and determine the binding state of each Rubisco complex, using the positions and orientations generated by STA analysis (31). These individual Rubisco are classified as “bound” (oligomer or polymer; nuclei are treated as polymers; this is necessary for mass balance in the theory) or “unbound” (monomer for nucleus size *n* = 2, or monomer plus dimer for nucleus size *n* = 3; discussed below). The bound species can be categorized by the number (*n*) of contiguous subunits comprising the polymer. A Rubisco identified as part of a polymer chain can be further processed with STA to isolate the Rubisco-Rubisco subunit interface (Fig. 1E-G). By using a numerical analysis rather than image classification, the analyst controls the parameters and corresponding error rates and can cleanly establish the polymer length and connectivity of each Rubisco; it is also computationally more efficient.

### Rubisco packing within α-CBs

The average volume of an untreated α-CB is 7 ± 2 ×10^−4^ μm^3^ corresponding to approximately a cube with 90 nm edge length or a sphere with 60 nm radius (Fig. 2A, Table 1). DTT treatment changed the average volume (Table 1), possibly through destabilization of the shell leading to selective loss of larger α-CBs (Fig. 2A, Table 1). Both chemical treatments resulted in populations with slightly higher Rubisco concentrations (45% for DTT treatment, 26% for diamide treatment, Table 1), presumably due to loss of less mechanically stable α-CBs during plunge-freezing. While the chemical treatments reduced the number of low-concentration data points available for polymerization analysis, enough remained to facilitate curve fits.

**Table 1.**
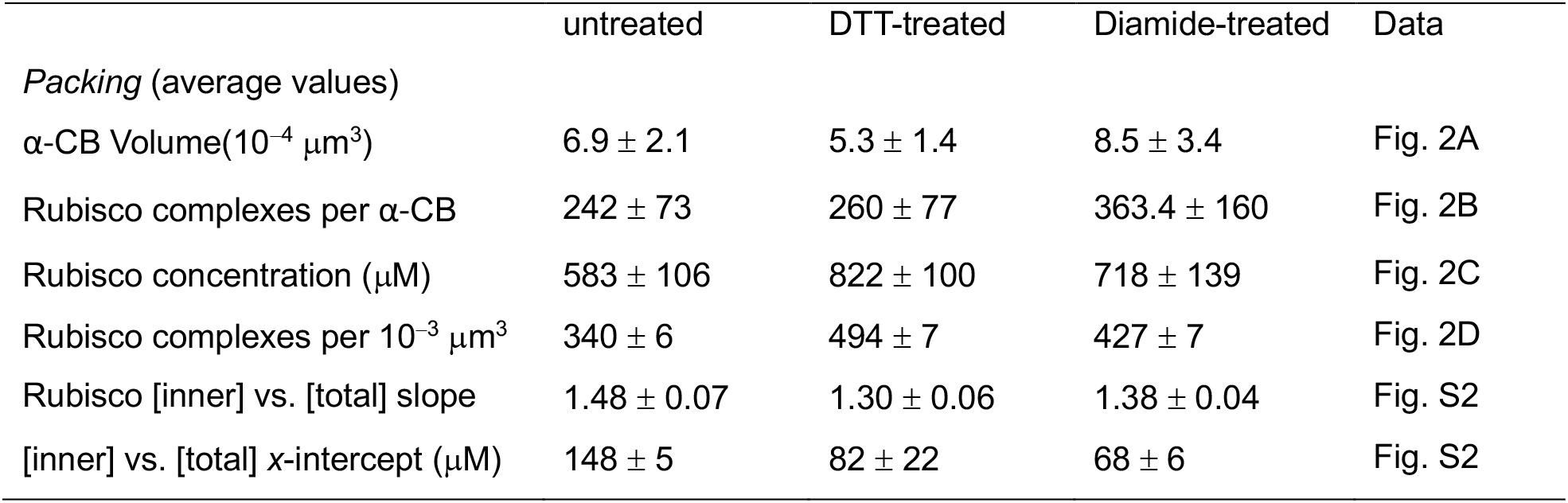
Rubisco packing properties inside α-CB.

| Table 1. Rubisco packing properties inside α-CB. | | | | |
| --- | --- | --- | --- | --- |
|  | untreated | DTT-treated | Diamide-treated | Data |
| *Packing (average values)* |  |  |  |  |
| α-CB Volume($10^{-4} \mu\text{m}^3$) | $6.9 \pm 2.1$ | $5.3 \pm 1.4$ | $8.5 \pm 3.4$ | Fig. 2A |
| Rubisco complexes per α-CB | $242 \pm 73$ | $260 \pm 77$ | $363.4 \pm 160$ | Fig. 2B |
| Rubisco concentration ($\mu\text{M}$) | $583 \pm 106$ | $822 \pm 100$ | $718 \pm 139$ | Fig. 2C |
| Rubisco complexes per $10^{-3} \mu\text{m}^3$ | $340 \pm 6$ | $494 \pm 7$ | $427 \pm 7$ | Fig. 2D |
| Rubisco [inner] vs. [total] slope | $1.48 \pm 0.07$ | $1.30 \pm 0.06$ | $1.38 \pm 0.04$ | Fig. S2 |
| [inner] vs. [total] x-intercept ($\mu\text{M}$) | $148 \pm 5$ | $82 \pm 22$ | $68 \pm 6$ | Fig. S2 |

**Figure 2.**
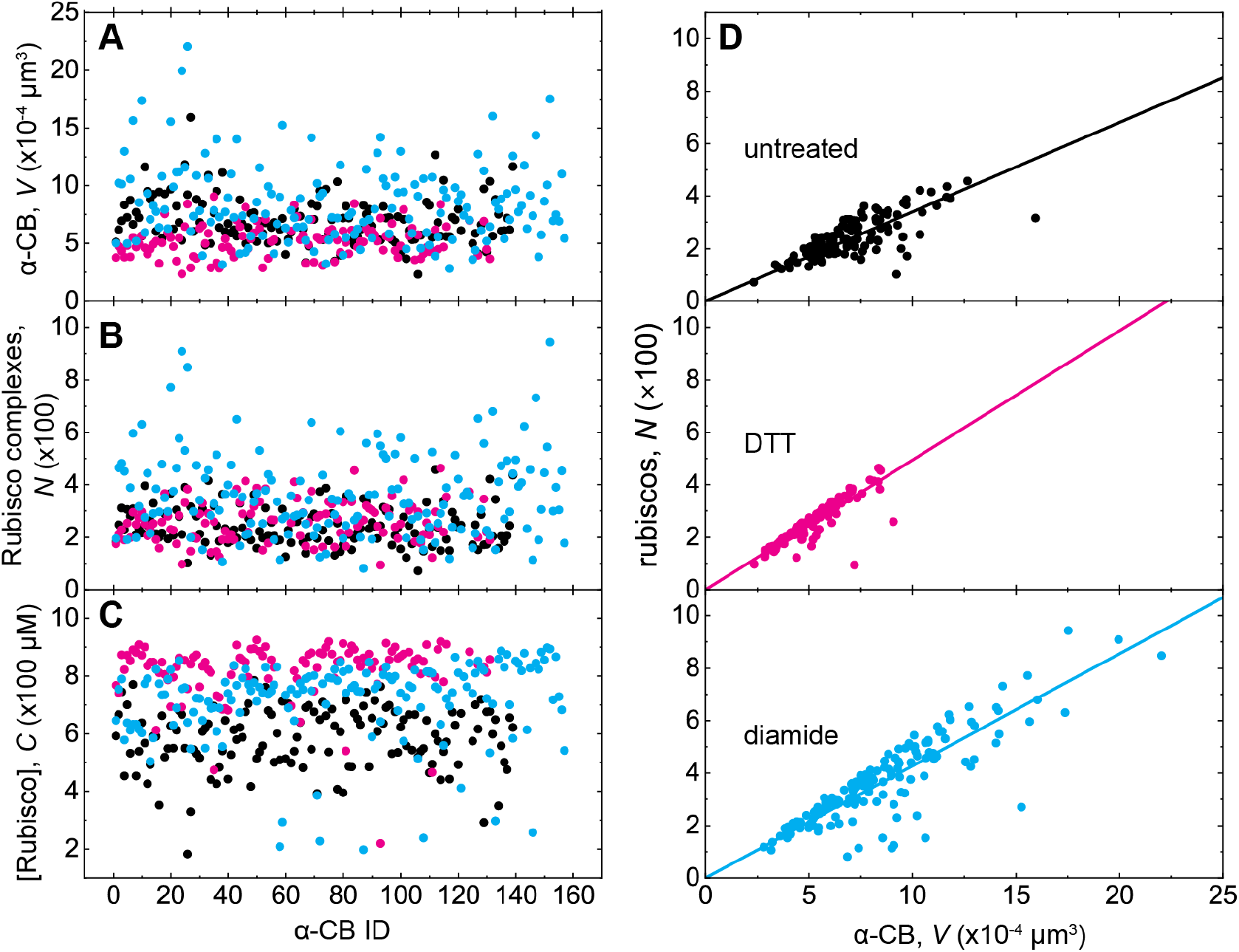
Rubisco packing in α-CBs. **A-C**. Comparisons of α-CB volume (**A**), number of Rubisco complexes in α-CB (**B**) and Rubisco concentration (*A*_*tot*_) in α-CBs (**C**) of untreated (black), DTT-treated (magenta), and diamide-treated (blue) samples. Data are summarized in Table 1. **D**. Correlation between the number of Rubisco complexes packed inside an α-CB vs. the volume.

The concentration of inner Rubiscos (defined as those not adjacent to the α-CB shell) scaled linearly with the total Rubisco concentration in untreated, DTT-treated, and diamide-treated samples (Fig. S2A-C). On average, inner Rubisco concentrations are about 1.4-fold higher than their total Rubisco concentrations for all three treated and untreated samples. The best fit lines to the inner Rubisco concentrations to total Rubisco concentrations for all three α-CB samples have a positive *x*-axis intercept (Fig. S2A-C, Table 1). This behavior indicates that inner Rubiscos are absent and all Rubisco is contained within the shell region at low total Rubisco concentrations, consistent with tomograms of lower-concentration α-CBs. This suggests that Rubisco near the shell may have different properties, interactions or other behaviors compared to the α-CB interior.

### Rubisco polymer chain length distribution within α-carboxysomes

Rubisco forms polymer chains in all α-CB populations imaged (Fig. 1B). Most of the polymers observed in the data are dimers, and the concentrations of longer chains decay with the chain length (Fig. 3). The Rubisco in dimers observed are not randomly contacting each other; rather, the interfaces observed between adjacent, continuous subunits in the Rubisco dimers are the same as those found in the longer polymers (Experimental section).

**Figure 3.**
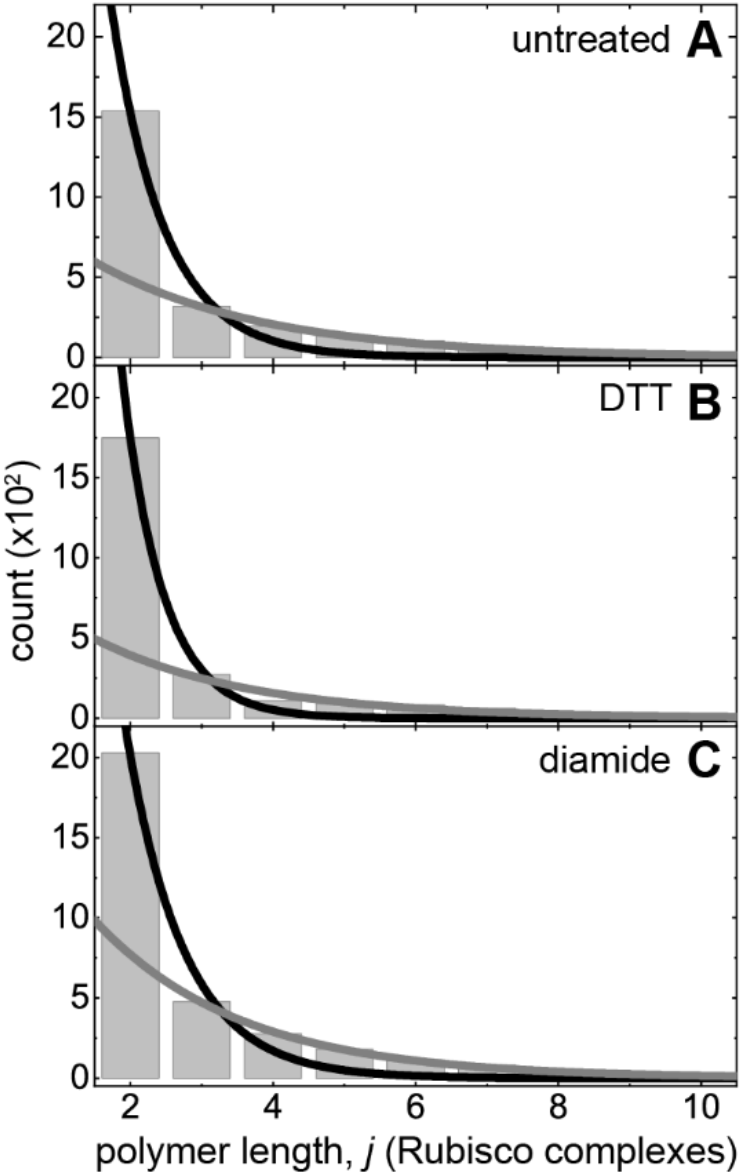
Length distribution of Rubisco fibrils in α-CBs. **A-C**. Histogram of polymer length for untreated, DTT-treated or diamide-treated. The continuous black lines are the best fits of the data to exponential decays including the dimer population (the first bar). The continuous gray lines (starting lower) are the best fits excluding dimers. Data available in Table 2.

The length distribution of an equilibrium polymer follows an exponential decay (32) (see Eq. 6 in Theory section). Histograms of Rubisco chain lengths inside all untreated, DTT treated and diamide treated α-CBs were fitted to an exponential (Fig. 3). The best fit to all the polymer lengths including dimers (*j* ≥ 2 Rubisco complexes; black smooth line) does not fit the data well, especially at longer polymer lengths. The observed dimer population deviates from the exponential decay defining the polymer population distribution, indicating that dimers are thermodynamically distinct from polymers of three or more Rubisco complexes. In contrast, the best fit to the polymer length distribution for trimers and longer polymers (*j* ≥ 3 Rubisco complexes; gray smooth line) fits the data well for all samples (Fig. 3). This exponential length distribution behavior indicates (Eq. 6) that the internal Rubisco interactions in polymers of three or more Rubisco complexes (*j* ≥ 3) are thermodynamically equivalent. We interpret this as an indication that the nucleus for Rubisco polymerization is three contiguous Rubisco complexes (*n* = 3).

**Table 2.** Rubisco fibril chain properties inside α-CB.

|  | untreated | DTT-treated | Diamide-treated | Data |
| --- | --- | --- | --- | --- |
| Distance between adjacent fibrils (nm) | $10.9 \pm 0.7$ | $10.3 \pm 0.5$ | $10.4 \pm 0.6$ | cryoET |
| Maximum fibril chain length, $m$ (Rubisco) | 10 | 10 | 11 | Fig. 3 |
| length decay const., $\kappa$ (+dimer; Rubisco<sup>-1</sup>) | $1.3 \pm 0.2$ | $1.8 \pm 0.2$ | $1.2 \pm 0.1$ | Fig. 3 |
| length decay const., $\kappa$ (−dimer; Rubisco<sup>-1</sup>) | $0.42 \pm 0.03$ | $0.46 \pm 0.08$ | $0.49 \pm 0.02$ | Fig. 3 |

A previous thermodynamic theory analysis (33) has shown that the chain length exponential decay constant (κ) is directly proportional to the total free energy of the polymer (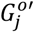 for a polymer composed of *j* Rubisco complexes) and the free energy change of the polymer 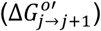 resulting from elongation by one subunit, in turn, can be linked to the polymerization constant (*K*_*pol*_; also referred to as the “critical concentration” for polymerization (34)) and the free monomer concentration in equilibrium with polymer (*A*_1_; Eq. 6 in the Theory section) according to:

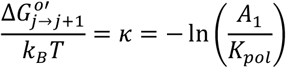

The free energy change 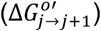 of the polymer when elongated by one subunit is given by:

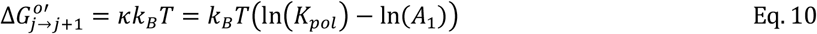

The free energy change per subunit extension 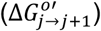 is constant for given *K*_*pol*_ and *A*_*tot*_, so the total value of 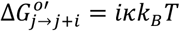 scales with the total number (*i*) of subunits incorporated into the polymer, including those resulting from the annealing of existing polymers.

The best fit to the Rubisco polymer (*j* ≥ 3) length distribution to an exponential (Fig. 3 and Table 2) yields 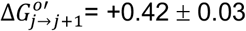, +0.46 ± 0.08, and +0.49 ± 0.02 *k*_*B*_*T* for untreated, DTT treated and diamide treated Rubisco complexes, indicating the Rubisco polymer chain internal stability is < *k*_*B*_*T* and comparable for all three samples. We emphasize that this free energy is distinct from the standard Gibbs free energy change (Δ*G*^0′^_*pol*_) associated with subunit addition to the filament end (discussed below).

### Rubisco polymerization mechanism within α-CBs

Because we determined the nucleus for Rubisco polymerization is a trimer (*n* = 3; Fig. 3), we treat Rubisco trimers and longer polymers as “bound” when plotting the concentration-dependence of polymer concentration (blue filled circles in Figs. 4 and S4 right panels). In this case, Rubisco dimers belong to the unbound, ‘monomer pool’ (light blue filled circles in Figs. 4 and S4 right panels). To extract the polymerization binding constants from this data, we can either fit the *A*_*tot*_ -dependence of bound Rubisco alone or globally fit both unbound and bound Rubisco concentrations together. Both fitting methods yield similar results, so we present only the global fits.

**Figure 4.**
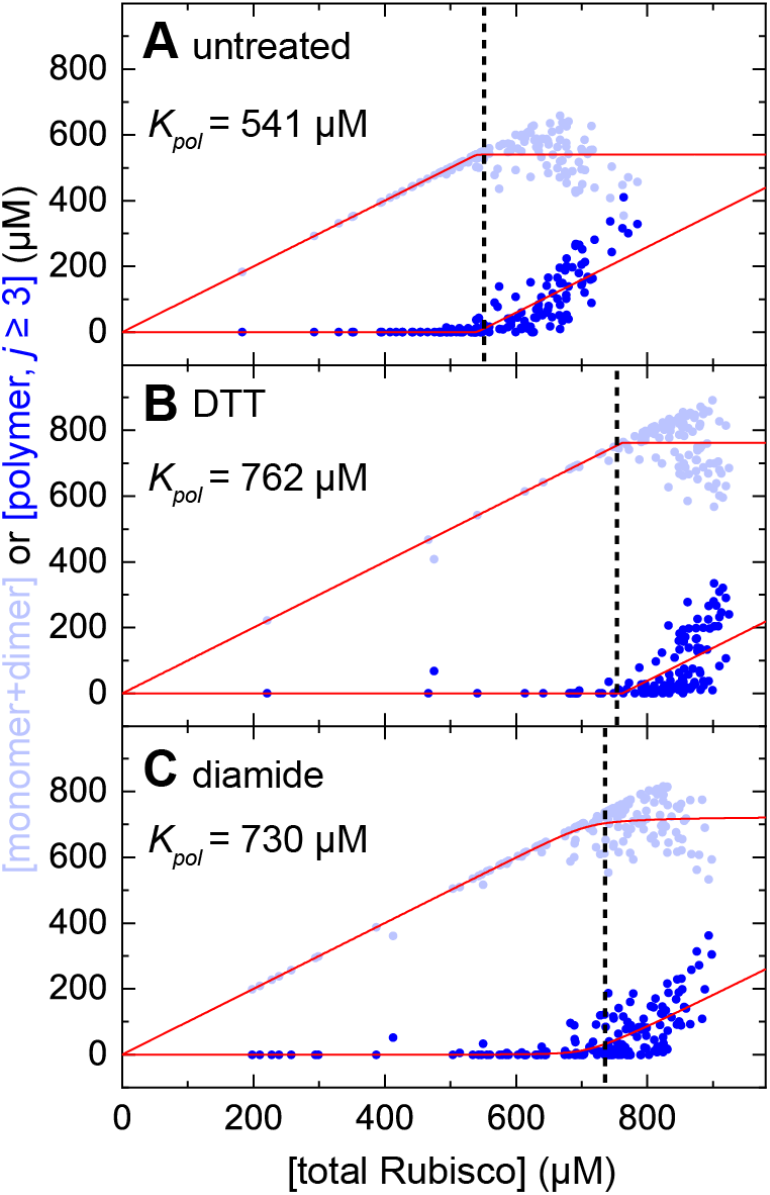
Rubisco polymerization analysis. **A-C:** Unbound (monomer+dimer, light blue filled circles) and bound (polymer of length *j* ≥ 3 Rubisco complexes; blue filled circles) Rubisco concentrations change as a function of the total Rubisco concentration inside untreated, DTT-treated, and diamide-treated α-CB samples. The smooth red lines through the corresponding data points represent the best global fits with implicit nucleation-polymerization equations for unbound (Eq. 3) and bound (Eq. 4) and the nucleation size of *n* = 3 held, respectively. The dashed vertical lines in all panels indicate the positions of *Kpol*.

The best global fit of the data to a length unlimited nucleation model (Eq. 3 and 4) with a trimer nucleus (*n* = 3) yields a *K*_*pol*_ = 541 ± 44 μM for untreated Rubisco, 762 ± 5 μM for DTT-treated Rubisco and 730 ± 170 μM for diamide-treated Rubisco polymers. These polymerization constants are weak (*K*_*pol*_ >500 μM), and associated with small standard Gibbs free energy changes for incorporation of a Rubisco subunit at a polymer end (Δ*G*^0′^_*pol*_) according to:

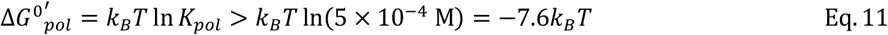

The nucleation binding constants (*K*_*nuc*_) are even weaker in all three cases, with values ranging from 10^9^ to 10^21^ μM (Fig. 4, Table 3).

**Table 3.**
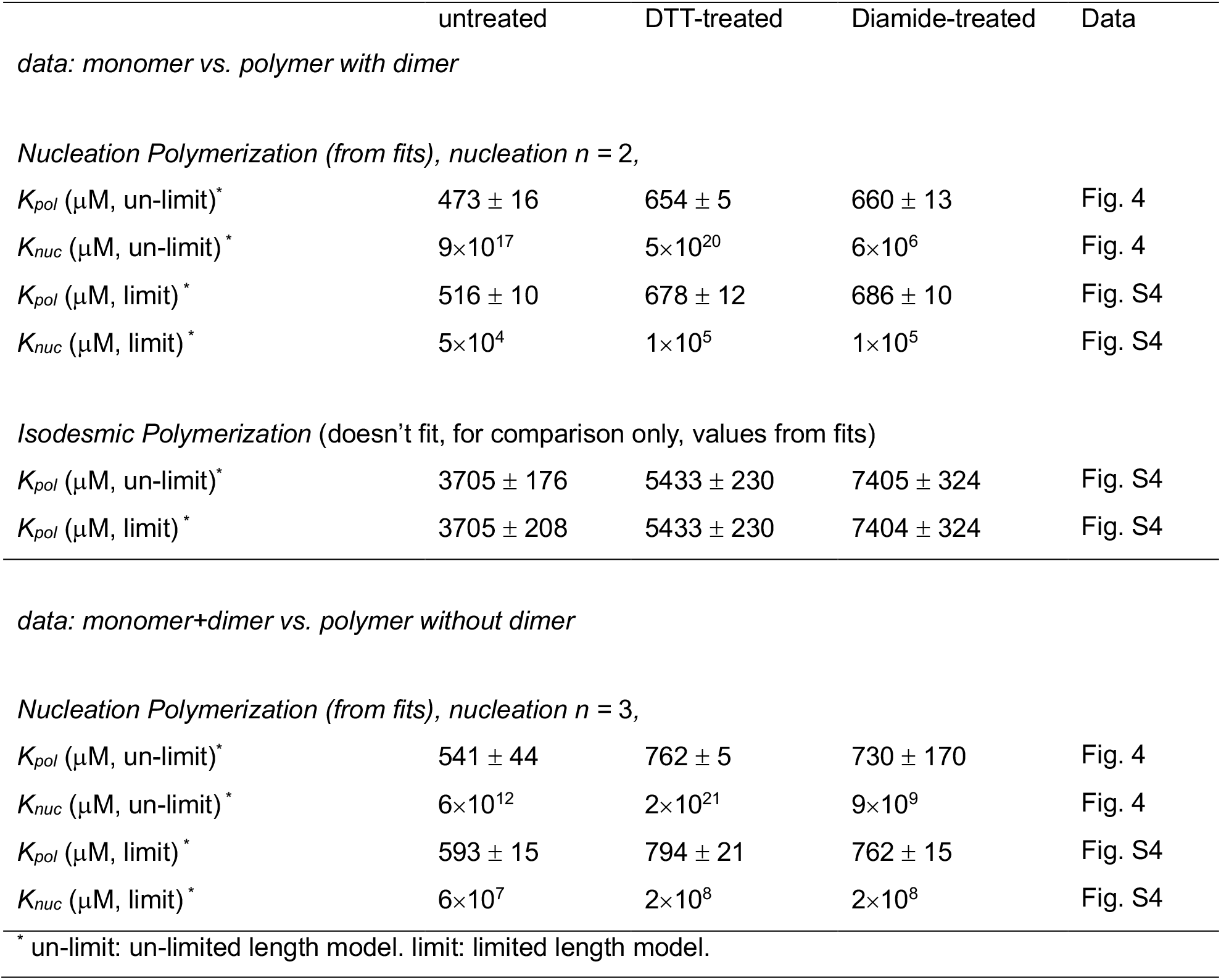
Rubisco polymerization properties inside α-CB.

For comparison, we analyzed the data with five other polymerization models: length-limited and length-unlimited isodesmic polymerization, length-limited and length-unlimited nucleated polymerization with a dimer nucleus (*n* = 2; Fig. S4A-C, left), and length-limited nucleated polymerization with a trimer nucleus (*n* = 3; Fig. S4D-F, right), together with length-unlimited *n* = 3 nucleated polymerization (Figs. 4 and S4D-F). In the case of isodesmic polymerization and dimer nucleation polymerization, unbound Rubisco complexes are treated as “monomers” (light gray filled circles), and bound Rubisco complexes are dimers and longer polymers (gray filled circles in Fig. S4A-C, left).

Isodesmic polymerization models do not fit the experimental data well for either length-limited or length-unlimited models (Fig. S4A-C, left). Dimer and trimer nucleation models with unlimited polymerization fit the corresponding data equally well (Fig. S4) and considerably better than with a length-limited constraint. The results of these analyses are summarized in Table 3. Fitting such polymerization data cannot determine the size of the nucleus (*n*) because the nucleus size was constrained when classifying oligomers as bound or unbound, thus, relating fit quality to nucleus size would be circular reasoning.

The best fits of the data to length-unlimited nucleated (*n* = 2 or 3) polymerization models yield very weak nucleation constant (*K*_*nuc*_) values with large uncertainties for all Rubisco samples (Table 3). The *K*_*nuc*_ values also vary by many orders of magnitude among the three samples. Despite the large uncertainties, the *K*_*nuc*_ values reflect and capture the sharpness of the monomer-polymer transition (i.e., inflection point) at the macromolecule critical concentration (Fig. S5). Abrupt transitions at the critical concentration indicate that nucleation (*K*_*nuc*_) is much weaker than subunit incorporation at a filament end (*K*_*pol*_). Self-assembling polymer systems with *K*_*pol*_ << *K*_*nuc*_ and abrupt transitions (i.e., actin filaments; (35)) are said to be “highly cooperative”. Thus, Rubisco polymerization is highly cooperative, despite the observed weak critical concentration. We note that despite the weak polymerization and large uncertainties in the collected data, the best fits can readily distinguish changes in the critical concentration for assembly under different conditions (Figs. 4 and S4).

## Discussion

### A novel method for evaluating protein-protein binding in STA

We demonstrate a novel approach to determine protein binding using volumetric imaging with STA, a technique typically used for protein structure determination (36). This approach leverages the intrinsic capability of STA to provide position and orientation information about proteins and other biological macromolecules, using these data to determine the relative concentrations of bound (i.e., polymerized, in a complex, etc.) and unbound particles in their biological context. This bottom-up method can be implemented as long as a protein can be clearly identified in both bound and unbound states for accurate classification. It is agnostic to concentration, allows experiments under native conditions, and does not require tagging or chemical modification of the target molecules. Concentrations can be calculated either from an enclosing barrier or directly from the particles’ occupied volume, either counting the particles alone or including molecular crowding as nonspecific electron density within the particle-occupied area (31).

Single-particle cryoEM has previously been used to estimate binding constants of purified proteins by resolving the structures of bound and unbound states. In structure-based drug design, cryoEM structures are used to select computational binding poses that correlate with compound potency, with binding energies validated experimentally by surface plasmon resonance or estimated computationally through energy minimization of predicted conformational changes upon substrate binding (37, 38). While this approach has shown promise for drug discovery, physiological relevance remains a concern: during plunge-freezing preparations, proteins are exposed to fluid shearing force, uncontrolled osmolarity increases, and surface interactions. Furthermore, the high-resolution structures required for atomic-level comparison preclude inclusion of the entire dataset, as less populated or heterogeneous states would not resolve. CryoET mitigates some of these concerns by imaging proteins within an enclosed system, such as a cell, microcompartment, or artificial lipid vesicle enclosing purified protein. These environments shield the interior from nonphysiological conditions during sample preparation, while supporting more complex three-dimensional analyses of protein organization and interactions.

Our model system, Rubisco in α-CBs, offers several advantages for the development of the method used here to identify protein assembly and determine binding and polymerization constants. Rubisco complexes are large and easily identifiable on a grid, and the presence of a visible shell with a defined volume provides secondary validation of occupied volume calculations for determining Rubisco concentration (31). The protein shell sequesters the interior chemically and mechanically over the sample preparation timescale, preserving the native conditions. Rubisco’s large size allows unambiguous particle identification in bound and unbound forms, and its marked conformational stability allows STA of the full population without the need for classification (30). Finally, the α-CB is relatively well characterized biochemically and relative abundances of the contained proteins known, providing validation for underlying calculations such as Rubisco concentration in the interior (39). Application of this new binding analysis methodology to other systems is certainly possible, with noise and error rates ultimately dependent upon the particle identification accuracy and final resolution of the STA dataset.

### Rubisco polymerization in α-carboxysomes

All the three Rubisco polymerization datasets yield an weak (large in value) polymerization constant *K*_*pol*_, (i.e., overall dissociation equilibrium constant for a monomer to bind either end of a polymer chain (40)), and an even weaker nucleation constant *K*_*nuc*_ (Table 3), consistent with the observed sharp transitions near the inflection point for all three samples (Fig. 4, S4 and S5). This supports a mechanism in which space confinement is a primary factor for Rubisco polymerization.

We tested four general models of macromolecule polymerization: isodesmic and nucleation models, both with and without spatial constraints. In unlimited space, the length of a self-assembling polymer chain is unconstrained and, in theory, can reach infinite lengths as the total subunit concentration approaches infinity. Under confinement, the polymer chain length is limited by the accessible space, geometry of the polymer chain, and the chain’s material properties.

The average diameter of an untreated α-CB is equivalent to the linear length of 10 or 11 Rubiscos complexes (see *Polymerization theory within a confined volume* in Theory of Protein Polymerization), suggesting the volume may limit the maximum allowed Rubisco chain length. The polymerization models must be modified to reflect this spatial constraint (Eq. 7). The compliance of the polymer can also contribute to the achievable length. For example, if Rubisco polymers are very compliant, such that they could bend and twist easily, the maximum allowed chain length could conceivably be longer than a rigid, linear polymer. In our data, the maximum Rubisco chain length observed is equivalent to the α-CB diameter (10 Rubisco complexes for untreated and DTT-treated and 11 for diamide-treated (Table 2)). This suggests that Rubisco polymers are rigid on the observed length scales and the chain length is limited by the accessible space within α-CBs.

All polymerization models tested here predict that the polymer chain length distribution at equilibrium follows an exponential decay (Eq. 6). The observed Rubisco chain length distribution shows that the dimer fractions deviate from the exponential distribution of the population, whereas trimers and longer polymers adhere to this exponential population distribution (Fig. 3). This suggests Rubisco polymerization follows a nucleation mechanism with nucleus size (*n*) of 3 Rubisco complexes.

Isodesmic polymerization models, in which nuclei are not required for polymer formation, do not fit data well (Fig. S4A-C). Nucleation models with either *n* = 2 or *n* = 3 with confinement yield reasonable fits to each corresponding data set, although the fits are worse than without confinement (Figs. 4 and S4) as indicated from the sharpness of the transition from unbound to bound species (i.e., the critical concentration inflection point).

The polymerization data were better fitted by the chain-length unlimited nucleation model (Figs. 4 and S4), suggesting that spatial confinement has limited effects on Rubisco polymerization. If the length limit was achieved before subunit packing, the observed filament length distribution would shift toward a uniform population with the spatially-limited length. Therefore, the Rubisco filament length distribution and polymerization analyses together favor a length-unlimited nucleation mechanism with a trimer (*n* =3) nucleus for Rubisco polymerization in α-CBs.

Previous α-CB assembly studies have returned conflicting results a tomography study observed α-CB-sized Rubisco condensates in the cytoplasm that the authors interpreted as assembly intermediates (41), suggesting that the Rubisco condensate would determine the α-CB size, while biochemical experiments indicate that the accessory protein CsoS2 is the primary determinant of α-CB shell diameter (42–44). We conclude that, while we cannot determine the details of α-CB assembly from these polymerization fits, we can successfully determine that the maximal Rubisco chain length is an intrinsic property of the condensate rather than imposed by the shell.

### Disproportional abundance of Rubisco dimers

Rubisco dimers are prevalent in condensates (Fig. 1A) as well as in α-CBs *in vitro* and *in situ*. Conversely, Rubisco trimers and longer polymers are found in low concentrations in solution, and are primarily in α-CBs(21).

We observed a large fraction of Rubisco dimers in α-CBs. This abundance of dimers is unexpected for a nucleus size (*n*) of 2 or 3, as oligomers shorter than the nucleus (*n* = 3) are typically unstable and populated at low concentrations, and oligomers equal to the nucleus should behave as polymers in the exponential length distribution (Fig. 3). The deviation of the dimer population from the exponential, polymer length distribution (Fig. 3), suggests that they are thermodynamically distinct from polymerized Rubisco fibrils.

The total dimer species is overpopulated, implying that it is more stable than any polymer species, and raising the possibility that a fraction of this dimer population is polymerization-incompetent. At least four cases can potentially account for this unexpected finding:

#### Case 1: Ambiguity identifying and quantifying the Rubisco dimer population

Two Rubisco monomers positioned in an adjacent manner could be incorrectly classified as a dimer, yielding an “overcounting” of the dimer population. (This would yield n=3 nucleus size, but with some dimers being monomers; see Figs. 4 and S4D-F.) However, the cryoET method employed here retains sufficient spatial and structural resolution to distinguish and confirm that the observed dimers are not randomly contacting each other but adopt the same orientation as adjacent, contiguous subunits in longer polymers. Furthermore, this packing effect should lead to similar “overcounting” of all observed oligomer and polymer species detected. We thus can eliminate miscounting as the source of the observed abundant dimer population.

#### Case 2. A population of Rubisco dimers are polymerization-incompetent and on the polymerization pathway

A mechanism in which two distinct Rubisco dimer conformers – one capable of polymerizing (*R*_2_) and one not (*R*_2_*) – are linked by a reversible and on-pathway isomerization equilibrium (*K*_*isom*_) can account for the observed overabundance of Rubisco dimers (Scheme 3a).

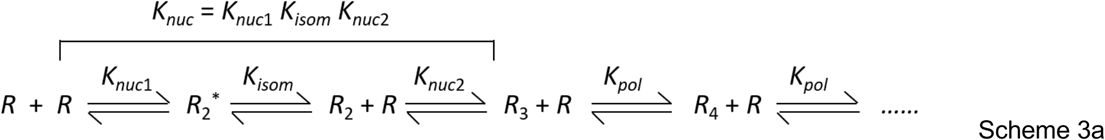

Given the resolution and orientation criteria for dimer classification (discussed in *Case 1*), the structural difference between the two dimer conformers would need to have a negligible effect on the Rubisco backbone, and likely be located at the subunit interface.

This polymerization pathway (Scheme 3a) defines a Rubisco trimer as the nucleus for polymerization (*n* = 3). The nucleation constant (*K*_*nuc*_) is defined here in a step-wise manner with binding constants *K*_*nuc*1_, *K*_*isom*_, and *K*_*nuc*2_, such that the overall nucleation equilibrium constant is given by the product of these three step-wise binding constants (*K*_*nuc*_ = *K*_*nuc*1_*K*_*isom*_*K*_*nuc*2_). This allows both the polymerization-incompetent dimers (*R*_2_*) and the nucleus (*R*_3_) to be expressed in terms of the polymerization-competent dimer (*R*_2_) concentration at equilibrium and the corresponding dissociation equilibrium constants (*K*_*isom*_ and *K*_*nuc*2_) according to [*R*_2_^∗^] = [*R*_2_]*K*_*isom*_ and [*R*_3_] = [*R*_2_][*R*]/*K*_*nuc*2_.

Accordingly, the ratio of the total dimer concentration (i.e., [*R*_2,*tot*_] = [*R*_2_*] + [*R*_2_]) over the nucleus trimer (*R*_3_) concentration is given by (Eq. 12):

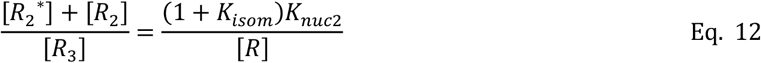

This relationship indicates the observed dimer population (*R*_2,*tot*_) will exceed that of the nucleus trimer (*R*_3_) when (1 + *K*_*isom*_)*K*_*nuc*2_ > [*R*] ∼ 500 μM (i.e., the Rubisco monomer concentration in equilibrium with polymer). Therefore, a weak *K*_*isom*_ (i.e., equilibrium largely favors polymerization-incompetent *R*_2_*) and/or weak *K*_*nuc*2_ (i.e., equilibrium favors *R*_2_ over *R*_3_) would generate an overpopulated dimer fraction.

In Scheme 3a, subunit addition to a dimer end (*K*_*nuc*2_) could be weaker or tighter than addition to a trimer end (*K*_*pol*_). If the ends of a Rubisco dimer and trimer are identical, as may be expected for a 1-D polymer, *K*_*nuc*2_ would be defined as *K*_*pol*_, *K*_*nuc*_ would be defined as *K*_*nuc*1_*K*_*isom*_, and the nucleus size would be a dimer (*R*_2_, *n* = 2), as detailed by (Scheme Sc3b):

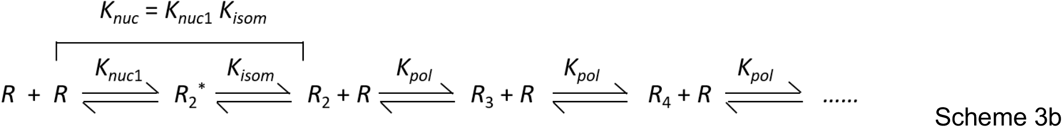

In this case, the nucleus *R*_2_ population adheres to the exponential, polymer-length distribution (Fig. 3), and the polymerization incompetent dimer (*R*_2_^*^) contributes to the over-counted dimer fraction. As with Scheme 3a, Eq. 12 with *K*_*nuc*2_ defined as *K*_*pol*_ predicts that the total observed dimer (*R*_2,*tot*_) will be overpopulated when *K*_*isom*_ is weak and largely favors *R*_2_^*^ such that (1 + *K*_*isom*_) >1.

#### Case 3. A population of Rubisco dimers is polymerization-incompetent and off-the polymerization pathway

A mechanism with two distinct Rubisco conformers (*R*_2_ and *R*_2_*), in which the incompetent dimer (*R*_2_^*^) is populated through a branched, off-pathway equilibrium can also potentially account for the observed population of Rubisco dimers. The non-productive, polymerization-incompetent dimers can form either through association of two Rubisco monomers (*K*_*dimer*_; Scheme 4a) or isomerization (*K*_*isom*_, Scheme 4b) of a polymerization-competent Rubisco dimer (*R*_2_) according to the following reaction pathways:

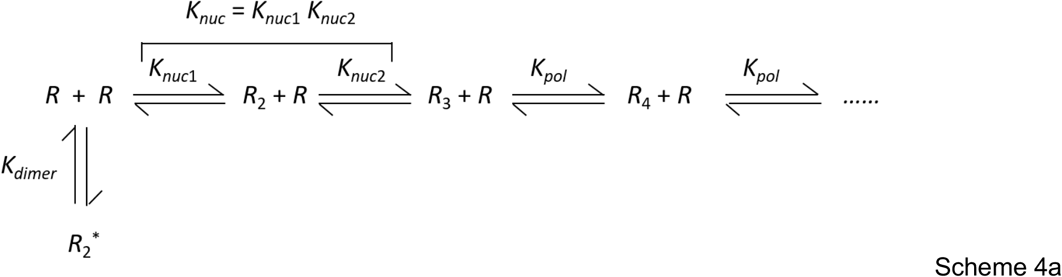

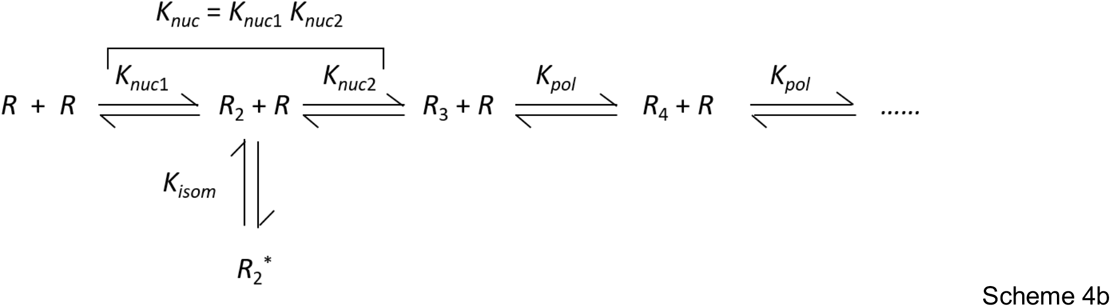

In Scheme 4a and 4b, if the nucleus size *n* = 3, the overall nucleation constant (*K*_*nuc*_) can be defined as *K*_*nuc*_ = [*R*]^3^/[*R*_3_] or in a stepwise manner as *K*_*nuc*_ = *K*_*nuc*1_*K*_*nuc*2_, with *K*_*nuc*1_ = [*R*]^2^/[*R*_2_]. The non-productive dimer concentration ([*R*_2_^*^]) is given by [*R*_2_^*^] = [*R*]^2^/*K*_*dimer*_ in Scheme 4a, and the following relationship holds (Eq. 13a):

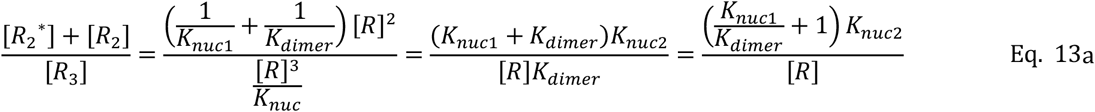

In Scheme 4b, [*R*_2_^*^] = [*R*_2_]/*K*_*isom*_ = [*R*]^2^/(*K*_*nuc*1_*K*_*isom*_), and the following relationship holds (Eq. 13b):

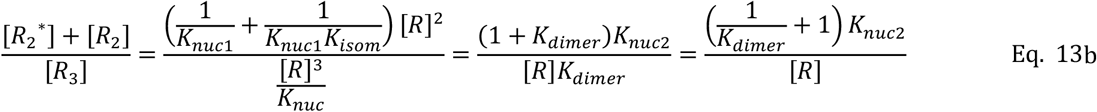

In both non-productive, off-pathway Rubisco dimer pathways, the observed dimer population will exceed the nucleus population (*R*_3_) in the polymer length distribution (Fig. 3) when *K*_*isom*_ or *K*_*dimer*_ favors *R*_2_* and/or *K*_*nuc*2_ is weak and favors *R*_2_ over *R*_3_.

As explained above in the case of an on-pathway, non-productive dimer (Scheme 3b), if the ends of a Rubisco dimer and trimer are identical, *K*_*nuc*2_ is defined as *K*_*pol*_, *K*_*nuc*_ is defined as *K*_*nuc*1_, and the nucleus size would be a dimer (*R*_2_, *n* = 2), according to:

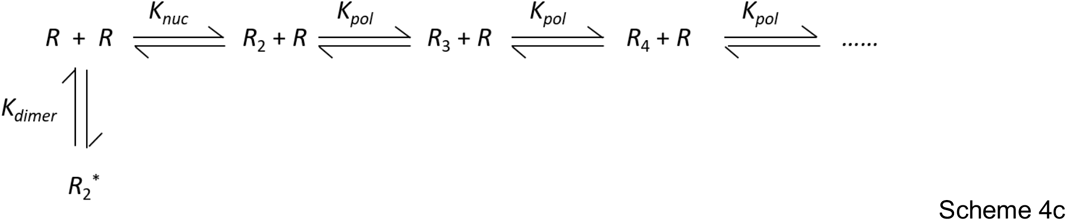

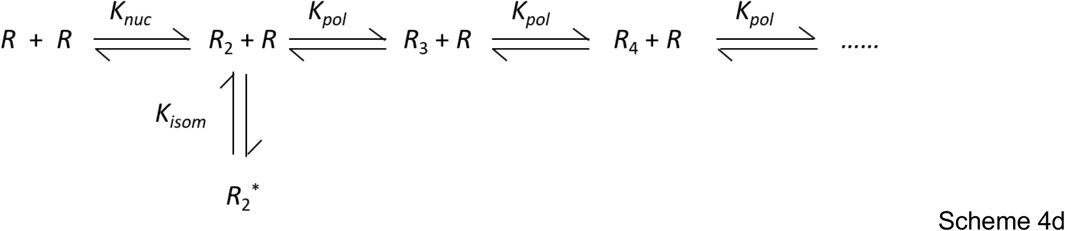

In these two cases, the total observed dimer population will be over counted when *K*_*nuc*_ is weaker than *K*_*dimer*_, thereby favoring *R*_2_*, or *K*_*isom*_ favors *R*_2_*, as above. As in *Case 2*, such an effect could not alter secondary or tertiary structure, or it would be visible in the structural biology analysis.

#### Case 4: A regulatory factor in α-carboxysomes binds and sequesters Rubisco dimers

An off-pathway, polymerization-incompetent dimer (*R*_2_^*^; *Case 3*) could potentially originate from the binding of a regulatory ligand (*L*), resulting in dimer sequestration. In this case, Schemes 4b and 4d become Scheme 5a and 5b, respectively, and all other relationships hold.

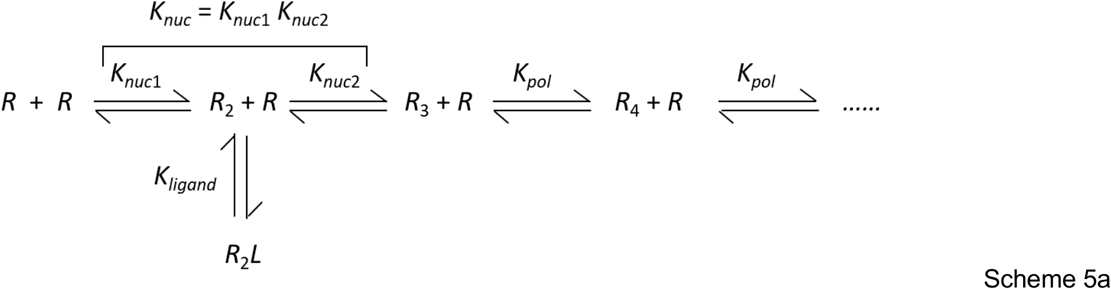

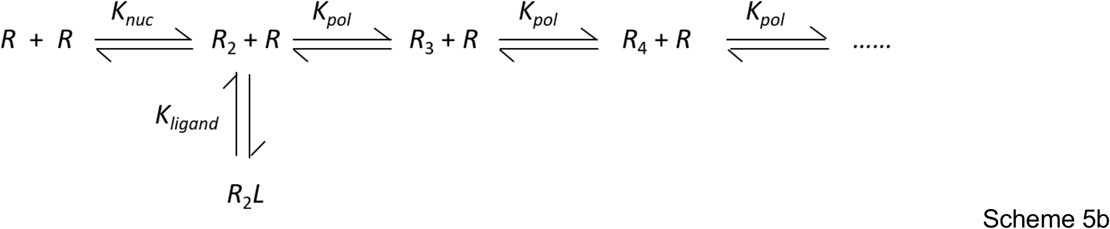

It is possible that a small-appearing ligand could be invisible in the STA structures if its occupancy was only once per complex, as the symmetry constraint in processing would diffuse its signal. While the STA was confirmed with no symmetry imposed (C1), a small-appearing ligand would have too small a signal to drive alignment.

#### Free energy changes associated with Rubisco polymerization

The total free energy of a polymer chain 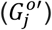 scales with the total number of subunits (Eq. 10) and thus increases when it elongates. The free energy change of a Rubisco fibril associated with incorporation of one subunit 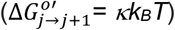 is < 0.5 *k*_*B*_*T* (Eq. 10) for all Rubisco samples evaluated here. A half *k*_*B*_*T* of energy change seems small since it is equal to the amount of the thermal energy associated with one degree of freedom, but it is >1200-fold larger than that associated with actin filament elongation 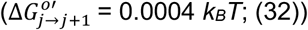. A polymer with larger 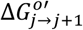 is thermodynamically and mechanically less stable and more susceptible to spontaneous fragmentation due to thermal and applied forces (45–47). Therefore, Rubisco polymer chains are considerably thermodynamically and mechanically less stable than actin filaments.

We clarify here for general readers that the two free energy changes for subunit incorporation at polymer ends discussed here, Δ*G*^0′^_*pol*_ vs 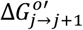, are not equivalent. The term 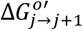 is the amount of free energy added to a polymer when it extends one subunit, while Δ*G*^0′^_*pol*_ is the free energy change for the overall reaction of a monomer binding to polymer end. Both are material properties, but 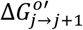 is an intrinsic constant, inversely scaling with polymers’ thermal and mechanical stabilities, while 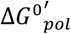 is an intrinsic constant reflecting the feasibility for monomer incorporation at a polymer end.

The (standard) Gibbs free energy change Δ*G*^0′^_*pol*_ is the negative change (loss) given by the difference between the Gibbs free energy (*G*^0′^) sum of a polymer with *j* subunits plus a free monomer (*G*^0′^_*j*_+ *G*^0′^_1_) and a polymer with *j* + 1 subunits (*G*^0′^_*j*+1_):

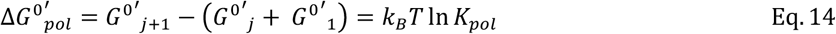

It reflects the free energy change associated with subunit incorporation at a filament end and is independent of the length of the elongating filament it is adding to. The polymerized subunit free energy is lower than that of a free monomer in solution, so subunit elongation is associated with a negative Gibbs free energy change.

In contrast, 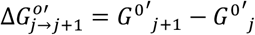 is defined as the (standard) free energy increase of the polymer itself when it elongates by one subunit (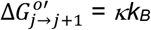, Eq 10). A longer polymer (of a given subunit type, such that the free energy change per subunit is constant) has a larger total free energy than a shorter one and is less therefore stable.

The two free energy changes refer to different reacting components are related according to the following, obtained by combining Eqs. 10 and 14:

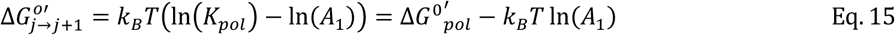

or

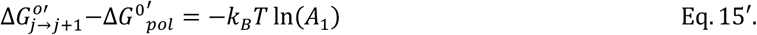

This relationship means the difference in the polymer free energy 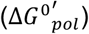 and the free energy (Δ*G*^0′^_*pol*_) for subunit incorporation is determined by the monomer concentration in equilibrium with polymer. The relationship between Δ*G*^0′^_*pol*_ and 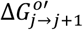 (Eq. 15 and 15’) can be independently obtained directly from the Gibbs free energy calculation (described in the Supplement).

### Relation of Rubisco chemical state and polymerization

One known chemical difference between cytoplasm and α-CBs is redox state: the cytoplasm is a reducing environment, while the interior of α-CBs is oxidized (28, 29). We therefore tested whether redox state could affect Rubisco polymerization. Using chemical treatments, we observed that the ordered lattices were often larger and more prevalent in oxidized samples compared to reduced samples, but with only subtle differences in *K*_*pol*_ (Figs. 4 and S4; Table 3). This finding emphasizes the importance of quantifying binding rather than using morphology as a guide: the decreased volume of the DTT-treated α-CBs likely affected this observation. α-CBs are purified without chemical reducing agents, and thus we anticipate that the untreated sample should be mildly oxidized as has been previously seen (48). One potential explanation is that the DTT treatment was not sufficient to fully shift the Rubisco polymerization, either due to the inability to revert side-chain oxygen additions or due to the relatively short treatment time tolerated by the α-CB shell.

If the Rubisco complexes exist in two populations within the α-CB (for example, reduced and oxidized), and if both populations polymerize and have different *K*_*pol*_ values within the observed *A*_*t*_ range, we would predict two inflection points in the data. Conversely, the sole inflection point observed here indicates that either only one polymerizing species is present, or that two polymerizing species exist but only one has a critical concentration within the observed range of *A*_*t*_. To test this, we introduced a floating parameter in the fitting program as the fraction of total Rubisco *A*_*t*_ that is either unable to polymerize at all or polymerizes beyond the observed *A*_*t*_ range. Fitting to all the three datasets resulted in this parameter being indistinguishable from zero, indicating that Rubisco polymerizes as a single species and most likely that redox state does not affect intrinsic polymerization capability in these experiments. We note that a case in which two species of Rubiscos exist but copolymerize is also consistent with the data.

For a system in which there is only one uniform polymerization species, α-CBs with the dense morphology (Figure 1B) should have Rubisco concentrations slightly below the *K*_*pol*_. However, a previous study of Rubisco lattice formation found that while the ordered morphology consistently had Rubisco concentrations above what we now know to be the *K*_*pol*_, the dense morphology covered a range of concentrations and could occur at Rubisco concentrations above this threshold(26).

Taken together, these findings suggest that changes to α-CB packing under different redox conditions are likely occurring through the action of accessary proteins, such as CsoS2, rather than through modification of the Rubisco itself (*Case 4* above).

### Conclusion

In summary, we have developed a novel tomography-based method for measuring polymerization in situ. In a model test of measuring Rubisco inside α-CBs, we were able to measure Rubisco *K*_*pol*_ and *K*_*nuc*_ constants inside an incompletely characterized physiological system with multiple protein species present, and at low affinities and high concentrations not typically amenable to in vitro study. We were able to determine that polymerization proceeds through a nucleus, and that redox-based effects on polymerization act through a secondary protein partner. This test case demonstrates the power of combining robust biophysical calculation and modeling with the *in situ* capabilities of cryo-electron tomography.

## Materials and Methods (abridged)

α-CBs were purified from wild-type *H. neapolitanus* as previous described (56).

### Negative stain imaging of condensate

Rubisco (0.5 μM) of L8S8 and the CsoS2 (2.7 μM) were mixed at room temperature for 1 hour. The sample was diluted 5x and Rubisco and CsoS2 condensates were visualized by TEM using a 2% (w/v) uranyl acetate stain. Imaging was performed on a JEOL 1200 EX transmission electron microscope.

### Sample Conditions and Tomography

The α-CB sample was diluted with BSA-coated 10 nm gold fiducials in a 4:1 ratio. Samples were treated with 5 mM diamide or DTT for one hour (57), then plunge-frozen onto glow-discharged C-flat 2/2-300 copper TEM grids. Imaging was performed at the Pacific Northwest Center for Cryo-Electron Microscopy on a Titan Krios G4 with a Gatan K3 camera and BioQuantum energy filter. Tomograms were collected in SerialEM according to Supplementary Table 2. Frame alignment, 2D image processing, and tilt alignment were performed in etomo; CTF estimation in ctffind4; and tomogram reconstruction in novaCTF following previously published pipelines (58–61).

### Subtomogram averaging

Particles were identified in SIRT-like filtered tomograms using a customized workflow designed to identify every Rubisco complex within each α-CB (26, 30). The general subtomogram averaging workflow was based upon a previous high-resolution approach originally derived from the Briggs group (9, 26). Subtomogram alignment and averaging was performed in Dynamo v.1.1.532, using the minimum number of iterations possible at each bin size and employing adaptive bandpass filtering throughout (62). Half-maps were sent to RELION5 for post-processing and final resolution calculations (63). Rigid body docking was performed in UCSF Chimera (64).

The Rubicso-Rubisco interface was generated by isolating all particles in trimers and averaging at the Rubisco-Rubisco interface. The model (Fig. 1E,F) showing the stacked Rubisco-Rubisco dimer was generated by fitting the monomer maps into the masked interface volume. The interface (Fig. 1G) is the STA volume generated by Dynamo.

### Data Analysis

We used a previously published Matlab script package to determine Rubisco binding states (31).

### Rubisco Filament Length Distribution Analysis

At equilibrium, the polymer length distribution is an exponential function of *A*_1_ (Eq. 6) and the unpolymerized monomer concentration *A*_1_ is function of *A*_*tot*_ (Eq. 1, 3, and 7) and increases with *A*_*tot*_, but *A*_1_ → *K*_*pol*_ as *A*_*tot*_ → ∞. For a nucleation polymerization, polymers appear only when *A*_*tot*_ > *K*_*pol*_, and in this region, *A*_1_ ∼ *K*_*pol*_ and is almost constant and changes little with *A*_*tot*_ (Fig. 4). Consequently, the polymer length distribution (Eq. 6) changes very little with *A*_*tot*_, and aggregating all polymer lengths across α-CBs with different *A*_*tot*_ (Fig 3) is acceptable for a length distribution analysis.

### Polymerization Analysis

The total Rubisco concentration-dependence of the monomer- and polymer-subunit concentrations were globally fitted to the chain length limited and unlimited isodesmic and cooperative nucleation-elongation polymerization models, (Eqs. 1-4 and 7) using Origin software, with the dimer treated as monomer or polymer depending on n = 2 or 3. A non-polymerizing fraction (non_poly) was included but was zero for all conditions. Implicit equations of the monomer concentration *A*_1_ were solved with a custom numerical routine based on a root-finding algorithm implementing the Newton-Raphson method combined with the Bisect method (65). Simulations were done with Matlab. For length-limited fits, m=10 (untreated, DTT) or 11 (diamide), with *A*_1_ ≤ *K*_*pol*_ but *K*_*pol*_ and *K*_*nuc*_ unconstrained.

## Data availability

Untreated carboxysome data were previously published and can be found at the Electron Microscopy Data Bank under accession code EMD-27654 and the Electron Microscopy public Image Archive under accession code EMPIAR-11125. The Rubisco unfiltered half-maps and full filtered map from both treated data sets have been deposited in the Electron Microscopy Data Bank under accession codes EMD-XXXXX and EMD-XXXXX. The complete set of subtomograms for each sample as well as exemplar reconstructed tomograms and frames have been deposited in the Electron Microscopy Public Image Archive under EMPIAR-XXXXX. All other data that support this study are available from the corresponding authors upon reasonable request.

## Acknowledgements

We thank Keith Meyers, Morgan Stephens, and Izabella Soswa for their assistance aligning tomograms and processing data for subtomogram averaging. Molecular graphics and analyses performed with UCSF Chimera (64).

## Authors’ contributions

L.A.M. and E.M.D.L.C. conceived and supervised the project. L.O. and D.F.S. provided the purified carboxysome samples and contributed images of the Rubisco/csoS2 condensate. L.A.M. prepared the samples for imaging and collected the tilt stacks. K.R., R.H.G., and L.A.M. processed the tomograms, conducted the STA, and carried out the biophysical processing of the particles. W.C. and E.M.D.L.C. established the mathematical theory and models, and interpreted the binding results. K.R., W.C., L.A.M., and E.M.D.L.C. wrote the manuscript. W.C. and K.R. prepared figures. K.R. finalized the structures and the EMDB/EMPIAR deposition. All authors contributed to the analysis and interpretation of the data.

